# Defining the Transcriptional Landscape during Cytomegalovirus Latency with Single-Cell RNA Sequencing

**DOI:** 10.1101/208603

**Authors:** Miri Shnayder, Aharon Nachshon, Benjamin Krishna, Emma Poole, Alina Boshkov, Amit Binyamin, Itay Maza, John Sinclair, Michal Schwartz, Noam Stern-Ginossar

**Author notes:** These authors contributed equally to this work (Miri Shnayder, Aharon Nachshon).

## Abstract

Primary infection with human cytomegalovirus (HCMV) results in a lifelong infection due to its ability to establish latent infection, one characterized viral reservoir being hematopoietic cells. Although reactivation from latency causes serious disease in immunocompromised individuals, our molecular understanding of latency is limited. Here, we delineate viral gene expression during natural HCMV persistent infection by analyzing the massive RNA-seq atlas generated by the Genotype-Tissue Expression (GTEx) project. This systematic analysis reveals that HCMV persistence *in-vivo* is prevalent in diverse tissues. Unexpectedly, we find only viral transcripts that resemble gene expression during stages of lytic infection with no evidence of any highly restricted latency-associated viral gene expression program. To further define the transcriptional landscape during HCMV latent infection, we also used single cell RNA-seq and a tractable experimental latency model. In contrast to current views on latency, we also find no evidence for a specific restricted latency-associated viral gene expression program. Instead, we reveal that latency-associated gene expression largely mirrors a late lytic viral program albeit at much lower levels of expression. Overall, our work has the potential to revolutionize our understanding of HCMV persistence and suggests that latency is governed mainly by quantitative changes, with a limited number of qualitative changes, in viral gene expression.

## Introduction

Human cytomegalovirus (HCMV) is a ubiquitous pathogen that, like all herpes viruses, can establish latent infection that persists for the lifetime of the host. In healthy individuals infection rarely causes any significant clinical symptoms due to a robust immune response (1, 2). In contrast, primary infection or reactivation from latency can result in serious and often life-threatening disease in immunocompromised individuals (3–5). Latent infection is, therefore, a key part of viral persistence and latently infected cells are a clear threat when the immune system is suppressed. Despite this, our molecular understanding of HCMV latency state is still limited.

HCMV is tightly restricted to humans, however in its host it has extremely wide cell tropism (6), and many kinds of cells can be productively infected, including fibroblasts, epithelial cells and smooth muscle cells (7). In contrast, latent infection was so far characterized only in cells of the early myeloid lineage, including CD34+ hematopoietic progenitor cells (HPCs) and CD14+ monocytes (8). It was further established that terminal differentiation of HPCs and CD14+ monocytes to dendritic cells or macrophages triggers virus reactivation from latency (9–13). This differentiation-dependent reactivation of latent virus is thought to be mediated by changes in post-translational modification of histones around the viral major immediate-early promoter (MIEP). These modifications drive the viral major immediate-early (IE) gene expression, resulting in reactivation of the full viral lytic gene program cascade and the production of infectious virions (11). Thus, the cellular environment is a key factor in determining the outcome of HCMV infection.

During productive lytic infection, HCMV expresses hundreds of transcripts and viral gene expression is divided into three waves of expression IE, early, and late (6, 14, 15). In contrast, the maintenance of viral genome in latently infected cells, is thought to be associated with expression of a relatively small number of latency-associated viral genes (16–21) in the general absence of IE gene expression. Due to their therapeutic potential, significant attention has been drawn to these few latency-associated viral gene products, but the full transcriptional program during latency remains unclear.

The earliest studies that looked for latency-associated gene expression identified a number of transcripts arising from the MIEP region of HCMV but no function was assigned to them (22–24). More systematic mapping of latency-associated transcripts was conducted with the emergence of microarray technology. Two studies detected a number of viral transcripts in experimentally latently infected myeloid progenitor cells (25, 26). The latent transcripts reported by these studies were not entirely overlapping, yet these findings were used as a guideline for targeted efforts to identify latent gene products. Interrogating the viral transcriptome in natural persistent infection is highly challenging since viral genomes are maintained in extremely few cells, at very low copy numbers, and viral genes are expected to be expressed in low levels. Nevertheless, subsequent work detected a number of these transcripts during natural latency (17, 18, 21), mainly using high sensitivity approaches such as nested PCR, building a short list of viral genes that is generally accepted to represent a distinct transcriptional profile during latent infection. These genes include UL138, UL81-82ast (LUNA), US28, as well as a splice variant of UL111A, which encodes a viral interleukin 10 (27–33).

More recently, RNA-seq was applied to map latency associated viral transcripts (34). This study revealed a wider viral gene expression profile that included two long non-coding RNAs (lncRNAs), RNA4.9 and RNA2.7 as well as the mRNAs encoding replication factors UL84 and UL44 (34).

Such genome-wide analyses are highly informative as they measure the expression of all transcripts in an unbiased manner. However, a major limitation is that they portray a mean expression in cell population, without reflecting intra-population heterogeneity. In the case of latent HCMV infection models this can be highly misleading since it is hard to exclude the possibility that a small, undesired population of cells, is undergoing lytic replication and thus can easily introduce “lytic noise”. This effect can be especially significant for viral genes that are highly expressed during lytic infection such as lncRNAs (15). Finally, the low frequency of natural latent cells is a major hurdle for global quantitative analysis of naturally latently infected cells.

To overcome the problem of scarcity of natural latent cells, we took advantage of the massive human RNA-seq atlas generated by the Genotype-Tissue Expression (GTEx) consortium (35). Through analysis of 435 billion RNA reads, we did not find any evidence for a restricted latency associated viral gene program. Instead, in several tissues we captured low-level expression of viral genes that resemble gene expression at late stages of lytic infection. Next, to directly explore viral gene expression in a controlled latently infected cell population we turned to the established myeloid lineage experimental systems. By using single cell RNA-seq (scRNA-seq) we unbiasedly characterize the HCMV latency program of both experimentally latently infected CD14+ monocytes and CD34+ HPCs, overcoming the impediment of cell population variability. Surprisingly, in contrast to the existing view in the field, we find no evidence for a specific latency associated viral gene expression program of only a few viral genes.

Instead, we reveal that in HCMV latency models, whilst there is little detectable IE expression, there is low-level expression of viral genes that strongly resembles the late lytic stage viral gene expression profile. Our analyses thus redefine HCMV latent gene expression program and suggest quantitative rather than qualitative changes that determine latency. Our work illustrates how new genomic technologies can be leveraged to reevaluate complex host-pathogen interactions.

## Results

### No evidence for a restricted latency-associated viral gene expression program in natural HCMV infection

We set out to characterize the full extent of viral transcription during latency in natural settings. The proportion of infected mononuclear cells in seropositive individuals was estimated at 1:10,000 to 1:25,000 with a copy number of 2-13 genomes per infected cell (36). Given that transcription of viral genes is expected to be low in these cells, immense amount of sequencing data is required to capture viral transcripts. We thus took advantage of the Genotype-Tissue Expression (GTEx) database, a comprehensive atlas containing massive RNA-seq data across human tissues that were obtained postmortem, from otherwise healthy individuals (35). We analyzed 9,416 RNA-seq samples from 549 individuals covering 31 tissues and containing more than 433 billion reads. This data-set (Fig. S1A) was aligned to the HCMV genome using strict criteria (material and methods). In 101 samples from 59 individuals we were able to recover HCMV reads (Fig. S1B).

When manually inspecting the reads that aligned to the HCMV genome we noticed that from 40 samples we obtained only reads that align to a well-defined, 229 bp region in the viral immediate early promoter (Fig. S1C). Although this region was previously associated with latency (22–24), we suspected these reads might originate from HCMV promoter-containing plasmid contamination. Indeed, when we analyzed HCMV genomes sequenced directly from clinical samples (37), we observed that in the majority of clinical samples, positions 175,493 and 175,494 in the genome (that fall in the MIEP region) contain cytosine or thymine in the first position and cytosine in the second position, whereas all the reads we identified in the GTEx samples matched the sequence of the HCMV promoter used in vectors which has adenosine and thymine in these positions (Fig. S1D). We therefore concluded that these reads may originate from a contamination and excluded them from further analysis.

For 51.7% of samples we had information about the CMV serostatus. Reassuringly, the number of samples that contained HCMV sequences and the number of HCMV reads were significantly higher in samples originating from seropositive individuals (Fig. 1A, Pval=0.0467 and Pval < 10^-55^ respectively, hypergeometric test). This is not due to differences in sequencing coverage as the read depth in samples from seronegative and seropositive individuals is similar. Further examination revealed that HCMV reads are found in 6 out of 2,210 seronegative samples, however all of them contained only one viral read per sample. Therefore this was used as a threshold and viral reads from samples containing less than two viral reads were filtered out in further analysis (data from all samples is summarized in Table S1).

**Figure 1:**
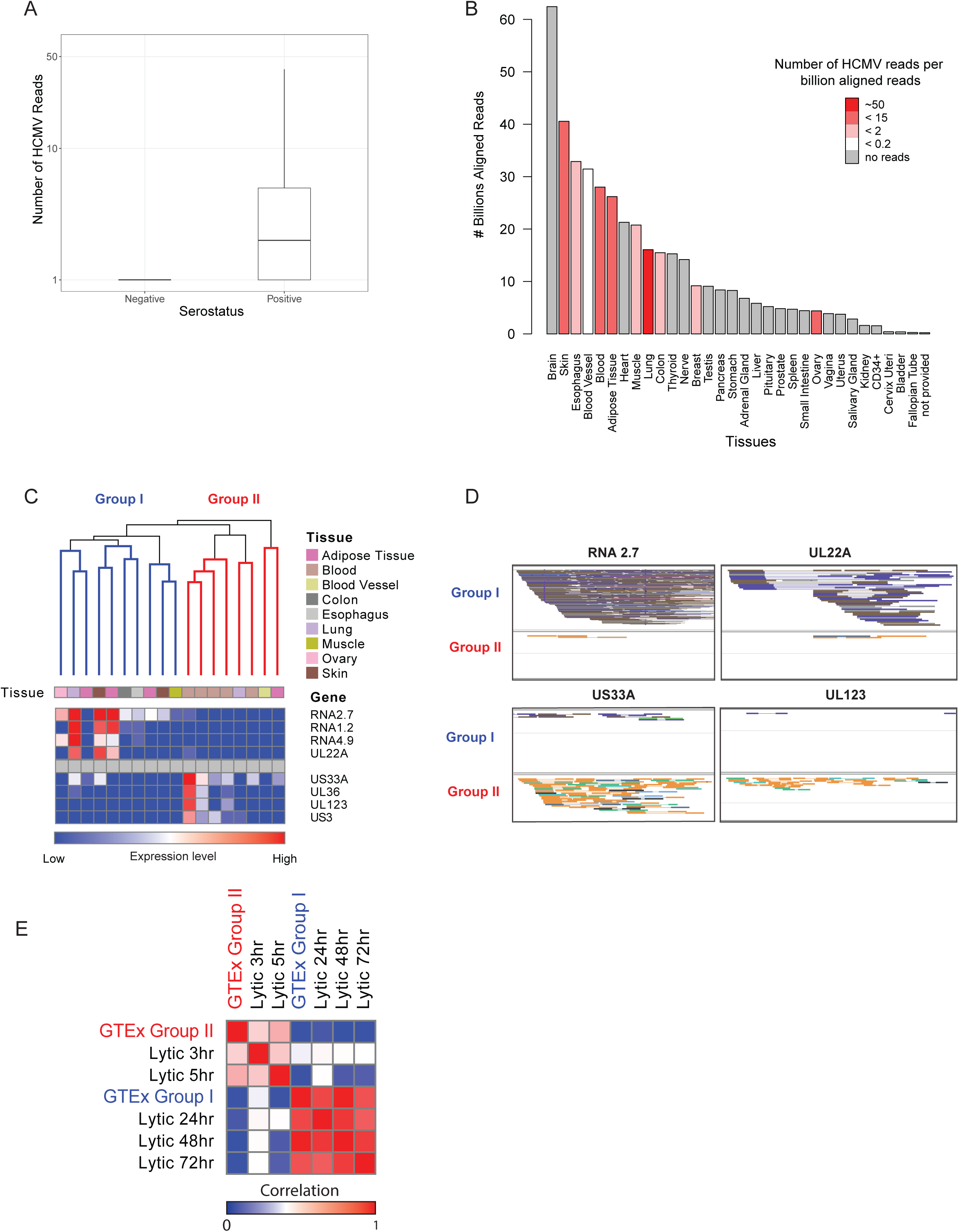
Viral gene expression during natural persistent infection. (A) Box plot showing number of HCMV reads per sample in HCMV seronegative and HCMV seropositive samples (B) Bar plot showing distribution of total sequenced reads in different tissues, color coding reflects the number of viral reads normalized to total number of sequenced reads in each tissue (Number of HCMV reads/10^9 total aligned reads). Viral reads from samples containing less than 2 viral reads were filtered out. Data for all samples was obtained from GTEx (35) except CD34+ data that were collected from 25 different NCBI GEO datasets (Table S2). (C) Hierarchical clustering of natural samples, with more than 4 HCMV reads, according to viral gene expression. The samples are portioned into 2 groups: group I and group II. Upper panel color coding indicates tissue origin of each sample. The heatmap in the lower panel shows expression level of representative differentially expressed genes in each sample. (D) Genome browser view showing aligned reads from samples assigned to group I or group II in genome regions coding for abundant genes in these groups. (E) Heatmap showing correlations between viral gene expression program from natural samples from both groups (I and II) and experimental lyticaly infected fibroblasts at different time points post infection.

Hematopoietic cells have been a primary focus of HCMV latency studies and HCMV genomes have been detected in HPCs and in additional cells throughout the myeloid lineage (38, 39). Consequently, the blood and the hematopoietic system are a major focus in research of HCMV persistence. The analysis of the GTEx database provides an exceptional opportunity to unbiasedly assess HCMV prevalence in various tissues. Interestingly analysis of the abundance of HCMV reads in different tissues revealed that ovaries, blood, adipose tissue and lung had the highest percentage of samples containing viral reads (Fig. S1E). The abundance of HCMV transcripts in these tissues was also the highest when analyzing the number of viral reads relative to the total reads that were sequenced from each of them (Fig. 1B). Since the GTEx database did not contain RNA-seq data from bone marrow where HPCs (which are the most established HCMV reservoir) reside, we performed RNA-seq on two HPC samples from HCMV positive individuals and surveyed additional 25 RNA-seq samples of HPCs from healthy individuals (Table S2). Although we analyzed over 1.5 billion aligned RNA-seq reads we did not detect any viral reads in these samples (Fig. 1B).

Next, we analyzed the viral gene expression as reflected by the HCMV reads we identified in natural samples, including in this analysis only samples that contained more than 4 HCMV reads. Hierarchical clustering revealed that the samples could be subdivided into two groups based on the pattern of viral gene expression (Fig. 1C). The first group (group I) was composed of samples that were dominated by transcripts that are the most highly expressed during late stage of lytic infection, e.g. RNA2.7, RNA4.9, RNA1.2 and UL22A (Fig. 1C and 1D). Indeed, when we compared the viral gene expression of these samples to RNA-seq data we collected along lytic infection of fibroblasts, we obtained high correlation to late stages of infection (R=0.97, Fig. 1E and Fig. S2A). This correlation suggests these viral reads that were identified in natural settings are similar to late stage lytic gene expression program.

The second group (group II) is composed of samples that express *bona fide* immediate early genes, e.g. UL123, US3 and UL36 as well as US33A which is the most highly expressed transcript early in infection (14), but had limited expression of transcripts that are abundant at late stage of lytic infection (Fig. 1C and 1D). Therefore, it is possible that these samples reflect the onset of viral reactivation, a state in which IE genes are transcribed but the full viral gene program is still suppressed. Supporting this notion, viral gene expression of these samples correlated best with lytically infected fibroblasts at 5 hours post infection (hpi) (R=0.55) (Fig. 1E and Fig. S2B). This IE expression positive state likely represent cells exiting from latency, consistent with the view that reactivation goes through a stage of IE gene activation. Since the tissues we analyzed were obtained postmortem, it is possible that postmortem-related physiological events led to HCMV reactivation and IE gene expression. To assess this hypothesis we inspected the time postmortem at which the tissue was collected (data is provided by GTEx (35)). Samples in group II were not enriched for long waiting time before tissue collection (Fig. S2C). In addition, there were no differences in the time interval of tissue collection between samples that contained HCMV reads and those that did not (Fig. S2D). These results suggest that the HCMV gene expression pattern we captured is largely independent of the trauma that occurred after death.

Unexpectedly, given the current view on cellular HCMV latency, we were not able to identify tissue samples that provide evidence for a restricted latency-associated viral gene expression program that differs from lytic viral gene expression.

### Single cell transcriptomic analysis of latently infected CD14+ monocytes

Although in natural samples we detected only low level viral gene expression pattern that resembles the lytic gene expression program, the cellular heterogeneity in these samples does not allow us to distinguish whether we are analyzing latently infected cells, or rare cells in which productive infection is taking place. Consequently, we next moved to characterize the viral transcriptome in experimental models of HCMV latency. Since these models rely on primary hematopoietic cells that may vary in their differentiation state and may also contain heterogeneous populations, we took advantage of the emergence of single cell RNA-seq (scRNA-seq) technologies (40, 41). This high-resolution profiling of single cell transcriptomes allowed us to delineate the nature of HCMV latency program in the best studied latent reservoir, hematopoietic cells.

Freshly isolated CD14+ human monocytes were infected with an HCMV TB40E strain containing an SV40 promoter driven GFP (TB40E-GFP) (42). This strain allows short-term detection of GFP-tagged latently infected cells, as in these cells GFP expression is efficiently detected at 2 dpi and then GFP signal gradually declines. Despite GFP levels in monocytes being much lower compared to those in lytic infection, the GFP expression allowed us to confirm that the majority of cells were indeed infected (Fig. S3A). To validate latent infection in our experimental settings, we analyzed by quantitative real-time PCR (qRT-PCR) the gene expression pattern of the well-studied latency associated gene, UL138 and of the immediate early gene, IE1 at 4 days post infection (dpi). Infected monocytes expressed relatively high level of UL138 while showing only trace level of IE1 transcript (Fig. 2A), thus manifesting the hallmark of latent infection (25, 27, 28, 33, 43, 44). Our latently infected monocytes release only trace amount of infectious virus, while differentiation of these infected monocytes into dendritic cells resulted in detectable IE expression as well as production of infectious virions (Fig. 2B and Fig. 2C). Thus we concluded that our CD14+ cells are latently infected.

**Figure 2:**
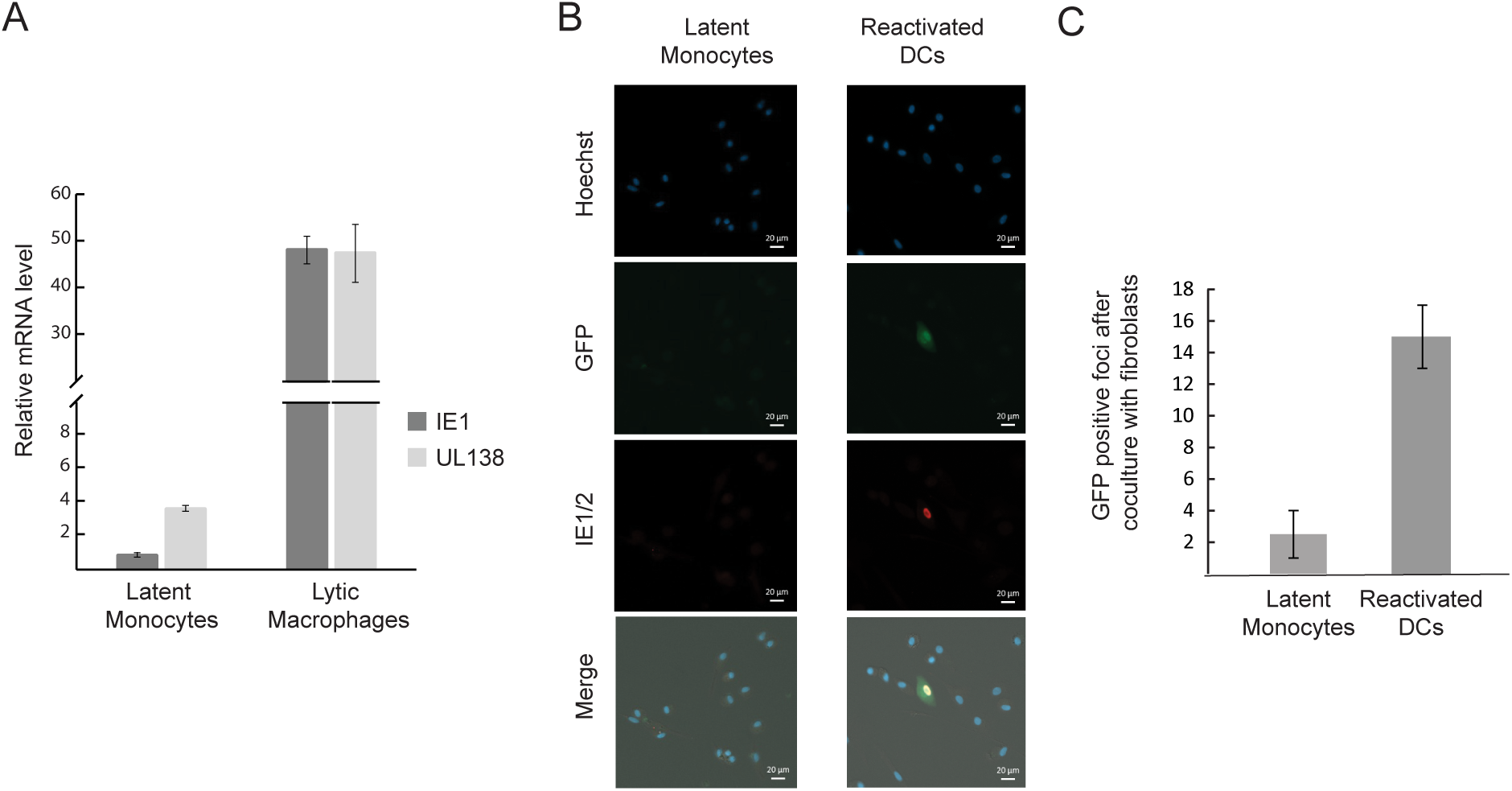
Establishment of HCMV latency in CD14+ monocytes. (A) Monocytes and monocyte-derived macrophages were infected with HCMV strain TB40E-GFP at an MOI of 5. RNA was collected at 4 days post infection (dpi) from the latent monocytes and 5 hours post infection (hpi) from lytic monocyte-derived macrophages and was analyzed by RT-qPCR for the transcript levels of UL138 and IE1. Expression was normalized to the human Anxa5 transcript. Means and error bars (showing standard deviations) represent three measurements. (B) Monocytes were latently infected with TB40E-GFP at an MOI of 5. Three dpi cells were either differentiated into dendritic cells (reactivated DCs) or left undifferentiated (latent monocytes) and 2 days post terminal differentiation reactivation was visualized by GFP and IE1/2 staining. Representative fields are presented. (C) Monocytes were latently infected with TB40E-GFP at an MOI of 5. At 3 dpi cells were either differentiated to dendritic cells (reactivated DCs) or left undifferentiated (latent monocytes). Two days post terminal differentiation cells were co-cultured with primary fibroblasts and GFP-positive plaques were counted. Number of positive plaques per 100,000 monocytes or monocyte-derived dendritic cells is presented. Cell number and viability were measured by Trypan blue staining prior to plating. Means and error bars (showing standard deviations) represent two experiments.

Next, HCMV infected CD14+ cells were single cell sorted without further selection at 3,4, 5, 6, 7 and 14 dpi and their transcript levels were measured using massively parallel 3’ scRNA-seq (MARS-seq) (45). Analysis of the entire transcriptome was performed on 3,655 CD14+ infected cells in which we could detect 15,812 genes out of which 171 were HCMV transcription units (see material and methods and Fig. S3B for distribution of reads and genes over the cell population). Projection of the cells using t-distributed stochastic neighbor embedding (t-SNE) analysis revealed that most of the cells constitute a large heterogeneous but continuous population and only a small group forms a distinct population (Fig. 3A). When we calculated the percentage of reads that align to the HCMV genome in each of the cells, it became evident that the small distinct population likely represents cells undergoing lytic infection as >10% of the reads in these cells originate from viral transcripts (Fig. 3A). Reassuringly, when performing the t-SNE analysis by using only cellular gene expression, we obtained the same structure, confirming we are looking at two different cell states (Fig. S4A). The small population represents a lytic cell population and the rest of the monocytes, which are the vast majority, exhibit very low to undetectable, diverse viral gene expression levels, indicating that they likely represent latently infected cells. This distribution, showing a clear separation between two groups of cells exhibiting very different levels of viral gene expression, confirms the purity of the single-cell isolation and the dominance of latent cells in the population of CD14+ infected cells (Fig. 3A).

**Figure 3:**
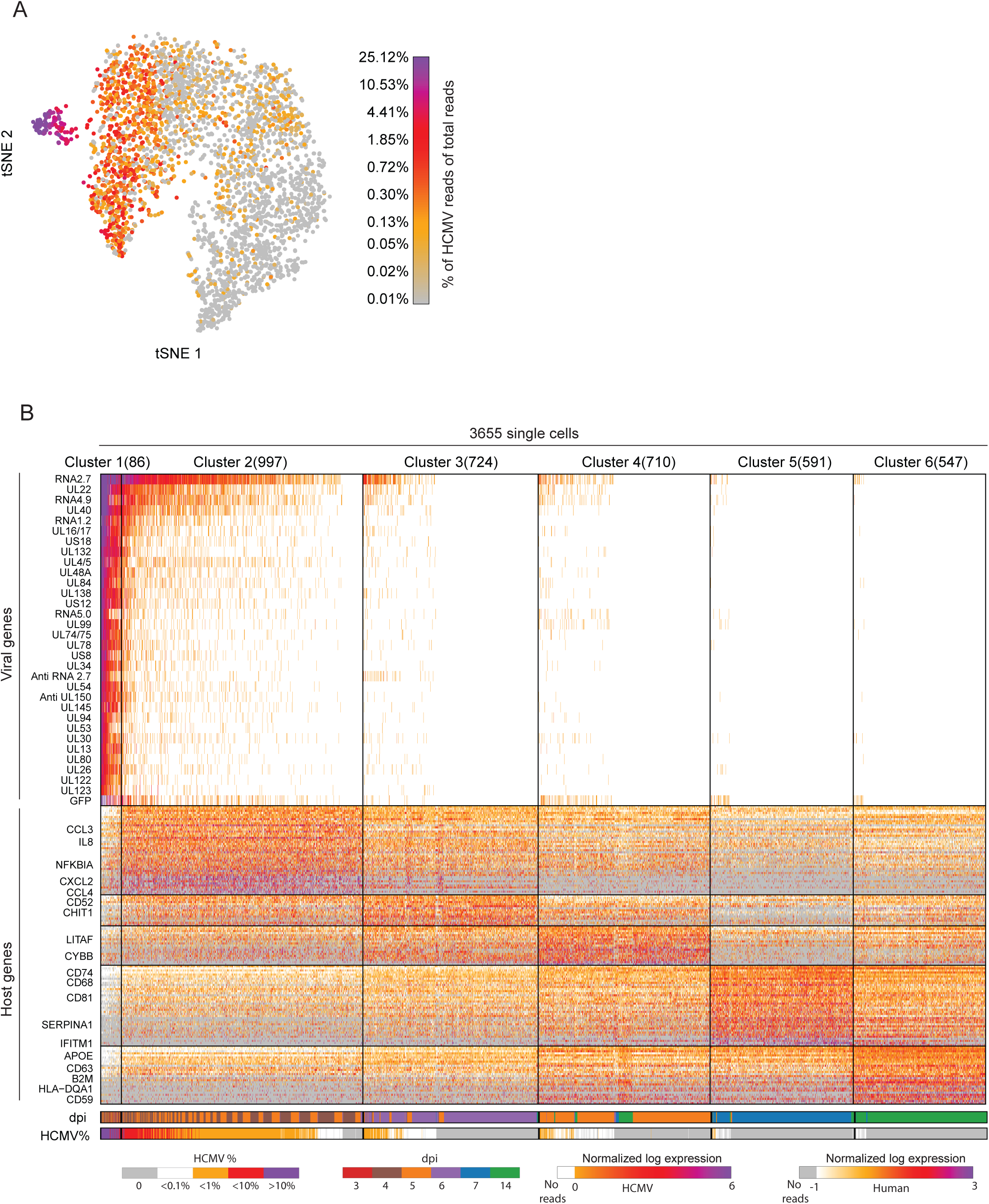
scRNA-seq analysis of latently infected CD14+ monocytes. Single cell RNA sequencing analysis of 3,655 cells from a cell population of latently infected monocytes. CD14+ monocytes were infected with HCMV (TB40E-GFP) and analyzed at 3, 4, 5, 6, 7 and 14 dpi. t-SNE plot of all 3,655 single cells based on host and viral gene expression. Color bar shows the percentage of viral reads from total reads per cell. (B) Heatmap showing clustering analysis of 3,655 single cells showing expression of 176 differential genes (32 viral, 144 cellular). Upper panel shows the most abundant viral genes, central panel indicates most differential host genes and lower panels indicate the percentage of viral reads from total reads and dpi for each cell. Cells are partitioned into 6 distinct clusters (C1-6) based on gene expression profiles and ordered by the relative abundance of viral reads, from high to low. Number of cells in each cluster is shown in parentheses next to the cluster number.

### HCMV latency associated gene expression in CD14+ monocytes and CD34+ HPCs resembles late lytic gene expression program

To assess the heterogeneity in HCMV latently infected monocytes, we combined the data from all 3,655 cells and clustered them on the basis of their host and viral gene expression profiles into 6 clusters (clustering method was previously described (46)) (Fig. 3B). Notably, also in this approach, the cells exhibiting high viral expression levels, presumably undergoing lytic infection, were clustered together and the most differential genes that were highly expressed in this cluster were almost exclusively viral genes (cluster 1, Fig. 3B, top panel). On the other hand, the rest of the cells exhibited very low levels of viral gene expression in varying degrees and the highly expressed differential genes in these five clusters were all cellular genes (Fig. 3B, lower panel and Table S3).

These clusters were consistent with the t-SNE analysis, with cluster 1 overlapping with the distinct population representing lytic cells (Fig. S4B). Indeed, by comparing the viral gene expression pattern of cells from this cluster to lytically infected monocyte-derived macrophages or fibroblasts we could confirm that they exhibit similar programs (Fig. S4C). Unexpectedly, although the lytic and latent cells represent two very separable cell states (Fig. S4A), latent cells from all clusters, show viral gene expression profile that is similar to the lytic cells (cluster 1), with the difference being in the *level* of viral gene expression but not in the *identity* of the viral genes (Fig. 4A). The only viral genes that their deviation from this correlation was statistically significant, and were relatively higher in latent cells, were the exogenous GFP (False discovery Rate (FDR)=7.10^-19^) which is driven by the strong SV40 promoter, the lncRNA, RNA2.7 (FDR<10^-100^), which is the most abundant transcript, and a transcript encoding for UL30 (FDR =6.10^-8^), a poorly characterized coding gene (15) (Table S4).

**Figure 4:**
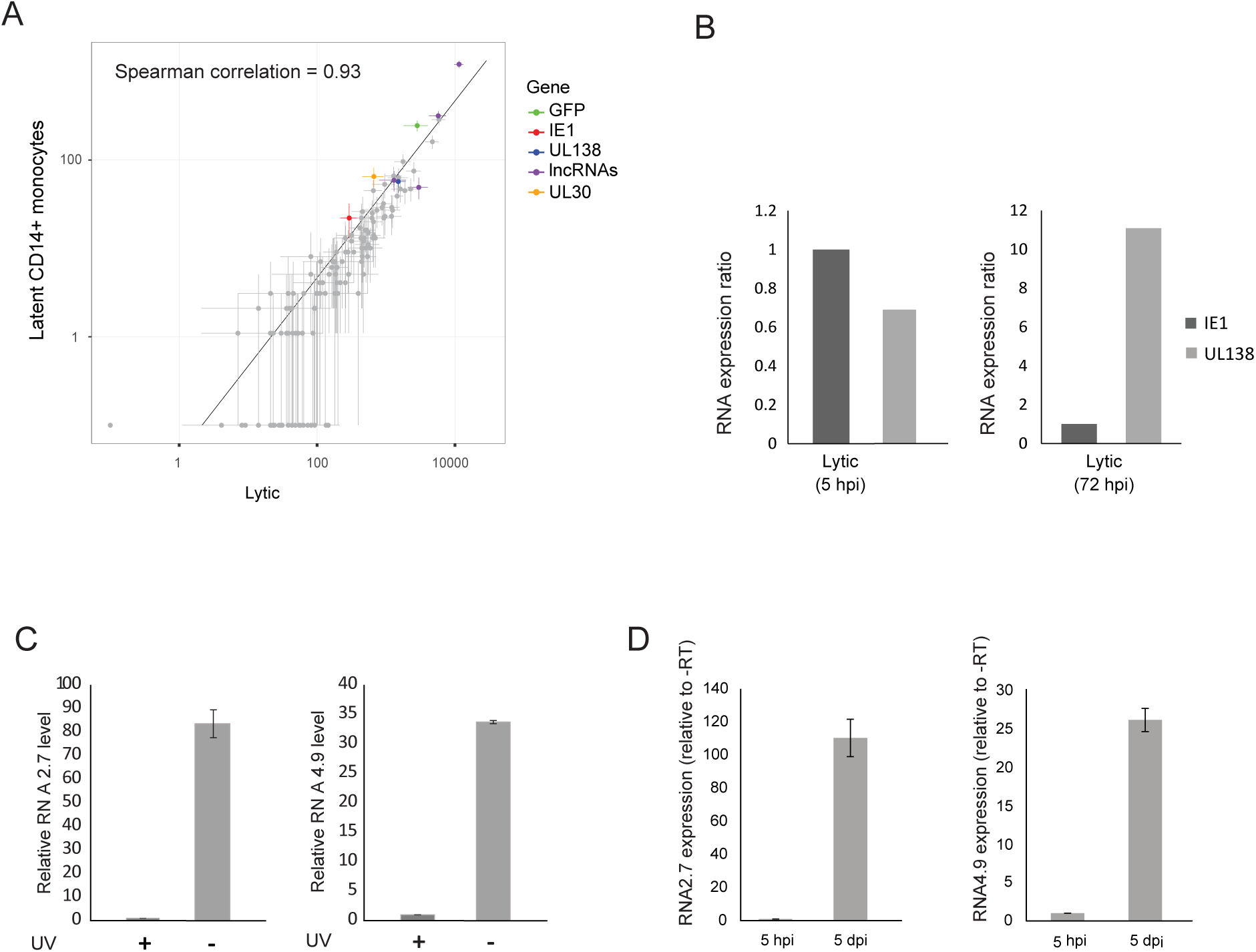
Transcriptional program in latently infected CD14+ monocytes. (A) Scatterplot showing read number of viral genes in latent monocytes (defined as cells in which the proportion of viral reads was below 0.5% of total reads) versus lytic cells (cells from cluster 1). Horizontal and vertical error bars indicate 95% non-parametric bootstrap confidence interval across cells. (B) Relative expression of IE1 and UL138 transcripts in RNA-seq data from lytic fibroblasts at 5 and 72 hpi. (C) Relative RNA expression level of viral RNA2.7 (left panel) and RNA4.9 (right panel) in monocytes infected with untreated or UV inactivated virus, measured by qRT-PCR at 5 dpi. A representative analysis of two independent experiments is shown. (D) RNA expression level of viral RNA2.7 (left panel) and RNA4.9 (right panel), relative to –RT samples, in infected monocytes, measured by qRT-PCR at 5 hours and 5 days post infection. Means and error bars (showing standard deviations) represent three measurements. A representative analysis of two independent experiments is shown.

We also examined whether viral gene expression program varies between the different populations of latently infected cells defined by the different clusters, by assessing the correlation between lytic cells (cluster 1) and each of the five other clusters. We found that viral gene expression profiles of all clusters were correlated with the lytic cells (Fig. S4D), with the correlation co-efficiency declining with the reduction in number of viral reads but only few viral genes were significantly higher in latent cells composing these clusters (table S5).

Importantly, by calculating the background noise in the single cell data (materials and methods), we confirmed that the results are not skewed by possible cross contamination in the single-cell data from the few lytic cells we have in our experiments (Fig. S5).

Overall, this analysis indicates that to large extent the viral gene expression program during experimental latency mirrors the viral gene expression program in late stage of lytic infection albeit expressed at much lower levels.

It is noteworthy that these unexpected results do not contradict previous analyses of latent cells, as we observe latent infection to be associated with overall low levels of viral gene expression and with high levels of UL138 relative to IE1. Importantly, this high UL138/IE1 ratio is also evident at late stages but not at early stages of lytic infection (Fig. 4B).

It was previously demonstrated that HCMV virions contain virus-encoded mRNAs (47, 48). To exclude the possibility that the transcripts we capture originate from input mRNAs that are carried in by virions, we infected CD14+ monocytes with untreated or UV-inactivated viruses and evaluated the levels of RNA2.7 and RNA4.9 at 5 dpi. The expression of both transcripts was over 30-fold lower in the cells infected with UV-inactivated virus compared to cells infected with untreated virus (Fig. 4C). In addition, viral transcripts levels at 5 hpi were much lower than at 5dpi (Fig. 4D), illustrating that the viral transcripts that we capture during latency result from de novo expression and are not the result of input mRNAs.

We next examined viral gene expression in experimentally infected CD34+ HPCs, which are another well-characterized site of latent HCMV infection (39, 49). CD34+ cells were infected with TB40E-GFP virus in the same manner as CD14+ monocytes, and used for generation of scRNA-libraries at 4 dpi. We initially used MARS-seq (45) to measure the transcriptome of infected HPCs, however out of 424 cells we sequenced, viral transcripts could be detected in only 12 cells (Table S6). We therefore moved to 10X genomics drop-seq platform that allows simultaneous analysis of thousands of cells. We analyzed the transcriptome of 7,634 experimentally infected HPCs in 366 of which we identified viral transcripts (see material and methods and Fig. S3C for distribution of reads and genes over the cell population). Projection of cells using t-SNE analysis revealed heterogeneous populations and cells that expressed viral transcripts were distributed throughout these populations (Fig. 5A). Analysis of the 366 cells that expressed viral transcripts revealed low expression levels and, as in CD14+ monocytes, viral gene expression in these cells resembled the expression pattern of late stage of lytic infection (Fig. 5B). Also here, there were only few transcripts that their deviation from this correlation was statistically significant, including RNA2.7 and UL30 (Table S7). Over all, our results show that during experimental latent infection there is no well-defined latency associated viral gene expression program, but rather these cells are characterized by low level expression of a program strongly resembling late lytic infection stages.

**Figure 5:**
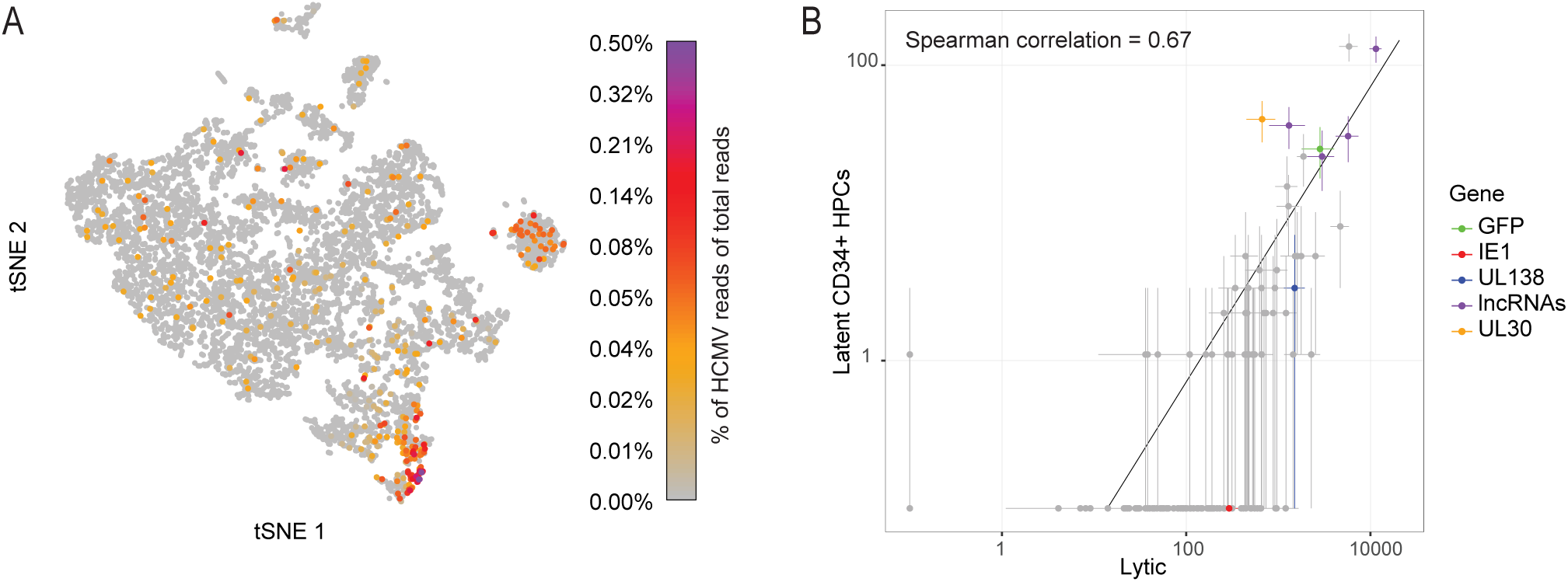
scRNA-seq analysis of latently infected CD34+ progenitor cells. Single cell RNA sequencing analysis of 7,634 cells randomly sampled from a cell population of latently infected HPCs. CD34+ HPCs were infected with HCMV (TB40E-GFP) and analyzed at 4 dpi (A) t-SNE projection of all 7,634 single cells based on host and viral gene expression. Color bar shows the level of viral gene expression as percentage of total reads per cell. (B) Scatter plot showing read number of all viral genes in the latently infected CD34+ progenitors versus lytic cells. Horizontal and vertical error bars indicate 95% non-parametric bootstrap confidence interval across cells.

## Discussion

Despite the clinical importance of HCMV latency, the mechanisms involved in viral genome maintenance and reactivation are poorly understood. An important step in deciphering these mechanisms is to characterize viral transcripts that are expressed during latent infection in an unambiguous manner. To address this challenge we examined HCMV infection by comprehensive analysis of RNA-seq data from diverse human tissues and further used scRNA-seq to analyze gene expression of latently infected CD14+ monocytes and CD34+ HPCs. Surprisingly, our measurements demonstrate that in both natural HCMV infection and in experimental latency models there is no evidence of a unique latency-associated gene expression program but instead we describe viral gene expression pattern that is similar to late stage of lytic infection at exceedingly low levels. Although these results are surprising given the prevalent notion that HCMV latency involves a specific gene expression program, evidence for broader viral gene expression was indicated in several previous genome-wide studies (25, 26, 34).

Examination of HCMV infection by analyzing viral gene expression in diverse human tissues did not reveal any restricted latency-associated viral gene expression program in the natural context. Instead we uncovered two patterns of gene expression; the first is composed of samples that contain viral transcripts that are abundant at late stage of lytic infection and the second is composed of samples with a restrictive gene expression pattern that includes mainly IE transcripts. The samples that contain late viral transcripts could reflect low-level expression that originates from few latent cells or the existence of scarce lytic cells in these tissues. The absence of IE gene expression implies that these reads may reflect a steady state rather than an ongoing lytic infection. Nevertheless, since cells expressing viral transcript are very rare, it is currently impossible to distinguish between these two scenarios.

The samples that contained mainly IE transcripts are fascinating as they may reflect a unique snap shot of viral gene expression during reactivation *in-vivo*, in natural human samples. Although we did not observe any difference in the time interval from death until these samples were collected, we cannot preclude the possibility that this restricted IE gene expression occurred postmortem or due to the associated trauma (50). Regardless of the conditions that initiated this restrictive IE gene expression, this state strongly suggests that *in-vivo* exit from latency goes through a phase in which IE genes are activated. The IE expression pattern we find was seen mostly in blood samples but not solely, suggesting that reactivation may occur in additional tissues, and at least in some stage is restricted to IE gene expression. The restrictive IE gene expression in these cells suggests that there is a threshold that needs to be crossed (perhaps the accumulation of enough IE proteins) before temporally controlled viral gene expression program can start. Indeed, this is entirely consistent with differentiation of CD34+ cells *ex-vivo* to immature DCs resulting in cells permissive for IE1 expression but not virus production (11) and with the detection of IE1 expression without infectious virus production in immature DCs isolated from healthy seropositive carriers (51).

Interestingly, our analysis of natural samples also suggest that HCMV persistence is widespread throughout the body, as we found viral gene expression in diverse human tissues. Previous studies have shown the presence of viral genomes in tissues outside the blood and hematopoietic system (Hendrix et al., 1997; Chen and Hudnall, 2006; Harkins et al., 2010; Gordon et al., 2017). Our data provide unbiased evidence for viral gene expression in various tissues and broader prevalence than was described before. The tissue that had the highest levels of viral transcripts was the lung which is consistent with recent results showing that HCMV DNA could be identified in the lung (Gordon et al., 2017), and in alveolar macrophages (9) and that HCMV reactivation is often manifested clinically as pneumonitis (57, 58). The cellular heterogeneity in tissue samples precludes any conclusion about the cellular sites of HCMV infection in these natural samples.

Our inability to detect a restricted latency-associated gene expression program in this systematic survey of natural samples, motivated us to examine the viral gene expression in the best studied latency experimental systems using single cell analysis. Notably, our results challenge the view of latency as being a specific viral-mediated program, and highlight rather a quantitative aspect of viral gene expression that is likely governed by the host cell. At the present sampling depth and coverage efficiency, our analysis of CD14+ cells can detect subpopulations of 0.3% (11–12 cells) or more. Therefore, although we cannot exclude the possibility that a small population of cells are in a different state and will harbor a different, more restricted, viral gene expression program, if such cells exist they would be rare.

The significant advantage of scRNA-seq, especially in the case of viral infection, is that we can exclude the possibility that the expression profile is skewed by a small group of cells. Importantly, in our work, the clustering approach allows us to validate that the viral gene expression profile is not related to viral expression levels as we see similar expression profiles even in the clusters that had significantly lower viral gene expression. Our analyses reveal differences in cellular gene expression that are associated with differences in the levels of viral gene expression. These differences could stem from variation in cell maturation state that restricts viral gene expression or alternatively they could reflect virally-induced changes in the host environment. Future work will help to distinguish between these two options.

The results we obtained for both CD14+ and CD34+ progenitors were qualitatively similar, however the relative levels of viral transcripts in CD34+ progenitors were significantly lower, suggesting that these cells are by nature more repressive. These results are in line with previous studies showing that MIEP is more repressed in CD34+ cells (59). Likewise, in natural latency we were unable to detect any viral transcripts by examining more than 1.5 billion RNA-seq reads from CD34+ cells. In contrast, by examining 3 billion RNA-seq reads from the blood we identified 378 viral reads from 18 samples. These results strongly suggest that viral gene expression is more restricted in CD34+ progenitors both in natural and in experimental settings and further illustrate that the host cell environment plays a major role in dictating the latency state.

An essential step in understanding HCMV latency is deciphering the importance of viral transcripts and proteins to latency maintenance and to the ability of the virus to reactivate. Based on the view that only a limited number of genes are expressed during HCMV latency, only several candidates for viral functions that may control HCMV latency have been studied. These include UL138 (27, 28), astUL81-82/LUNA (30, 44), UL111A/LAcmvIL-10 (29, 31) and US28 (32, 33). Despite the lack of a restricted latency expression program, our results do not undermine the importance of these factors to HCMV latency, rather add many additional candidate genes. Two appealing candidates are RNA2.7 and UL30. RNA2.7 is the most abundant transcript in both lytic and latent cells, but in our measurements RNA2.7 relative expression in latent cells was constantly higher than expected when comparing to the lytic profile. RNA2.7 was demonstrated to protect infected cells from mitochondria-induced cell death (60), but its role in latency was never tested. UL30 transcript was suggested to encode for *UL30A*, which is conserved among primate cytomegaloviruses, and expressed from a nonconventional initiation codon (ACG) (14, 15) but its functional role was never studied. Future work will have to delineate the importance of the different transcripts we detected to regulating latency.

Overall, our experiments and analyses challenge the dogma that all herpesviruses express a restricted latency-associated program and suggest that HCMV latency is characterized by a quantitative shift instead of qualitative change in gene expression. Although the relevance of these transcripts to latency should be further studied, these findings provide a new context for deciphering virus-host interactions underlying HCMV lifelong persistence.

## Acknowledgments

We thank Yosef Shaul, Schraga Schwartz, Igor Ulitsky, Rotem Sorek, Ian Mohr and Stern-ginossar lab members for critical reading of the manuscript. We thank Eain A. Murphy for the TB40E-GFP virus strain. We thank Elad Chomsky, Yaara Arkin, Hadas Keren-Shaul and Efrat Hagai for technical assistance. This research was supported by the EU-FP7-PEOPLE career integration grant, the Israeli Science Foundation (1073/14), Infect-ERA (TANKACY) and the European Research Council starting grant (StG-2014-638142). NS-G is incumbent of the skirball career development chair in new scientist.

## Supplementary tables

Table S1: Summary of HCMV reads in GTEx samples Columns indicate: Sample ID, subject ID, HCMV Sero Status, number of reads (in millions), number of aligned reads (in millions), number of HCMV reads, number of HCMV reads not including UL126. Columns I to end indicate the number of reds for each indicated gene.

Table S2: analysis of publicly available CD34+ RNA-seq datasets Columns indicate: Data set ID, Sample file ID, Cell type, number of reads in indicated sample, number of aligned reads.

Table S3: Differential genes in latently infected monocytes Columns indicate: Gene annotation, the cluster with the highest expression of the indicated gene, expression level of the indicated gene in the cluster where it is most highly expressed (relative to the expression of all other genes in the same cluster), number of reads for the indicated gene in clusters 1-6, total number of reads for the indicated gene across all clusters.

Table S4: Transcripts enriched in latent CD14+ monocytes Columns indicate: gene annotation, number of reads in lytic cells (cluster 1), number of viral reads in latent cells (cells in which less than 0.5% of the reads originated from the virus), mean and SD of the number of reads for an indicated gene in the latent cells according to bootstrap analysis (see material and methods), Z-score, false discovery rate (FDR).

Table S5: Transcripts enriched in latent cells from clusters 2-6 compared to lytic cells Columns indicate: gene annotation, number of reads in cluster 1, number of reads in the indicated cluster, mean and SD of the number of reads for each specified gene in the indicated cluster according to bootstrap analysis (see material and methods), Z-score, false discovery rate (FDR).

Table S6: Viral genes detected in latently infected CD34+ HPCs by MARS-seq analysis The number of reads identified for each of the detected viral genes, in each of the cells. Cell sum-indicates the total number of viral reads per cell. Gene Sum-indicates the total number of reads detected for each viral gene.

Table S7: Transcripts enriched in latent CD34+ HPCs Columns indicate: gene annotation, number of reads in lytic cells (cluster 1 in CD14+ analysis), number of viral reads in infected HPCs, mean and SD of the number of reads for an indicated gene in latent CD34+ HPCs according to bootstrap analysis (see material and methods), Z-score, false discovery rate (FDR).

## Materials and Methods

### Cells and virus stocks

Primary CD14+ monocytes were isolated from fresh venous blood, obtained from healthy donors, using Lymphoprep (Stemcell Technologies) density gradient followed by magnetic activated cell sorting with CD14+ magnetic beads (Miltenyi Biotec). Cryopreserved Bone Marrow CD34+ Cells were obtained from Lonza. Alternatively, fresh CD34+ cells were purified from umbilical cord blood of healthy donors. Isolation was done using Lymphoprep (Stemcell Technologies) density gradient followed by magnetic activated cell sorting with CD34+ magnetic beads (Miltenyi Biotec). CD34+ and CD14+ cells were cultured in X-Vivo15 media (Lonza) supplemented with 2.25mM L-glutamine at 37⁰C in 5% CO2 (61).

Human foreskin fibroblasts (HFF) (ATCC CRL-1634) and retinal pigmented epithelial cells (RPE-1) (ATCC CRL-4000) were maintained in DMEM with 10% fetal bovine serum (FBS), 2mM L-glutamine, and 100 units/ml penicillin and streptomycin (Beit-Haemek, Israel).

The bacterial artificial chromosome (BAC)-containing the clinical strain TB40E with an SV40-GFP tag (TB40E-GFP) was described previously (62, 63). This strain lacks the US2-US6 region, and therefore these genes were not included in our analysis. Virus was propagated by electroporation of infectious BAC DNA into HFF cells using the Amaxa P2 4D-Nucleofector kit (Lonza) according to the manufacturer’s instructions. Viral stocks were concentrated by ultracentrifugation at 70000xg, 4⁰C for 40 minutes. Infectious virus yields were assayed on RPE-1 cells.

### Infection and reactivation procedures

For experimental latent infection models, CD14+ monocytes and CD34+ HPCs were infected with HCMV strain TB40E-GFP at MOI of 5. Cells were incubated with the virus for 3 hours, washed and supplemented with fresh media. To assess infection efficiency, a sample of the infected cell population was FACS analyzed for GFP expression at 2 dpi. For single cell experiments cells were isolated without further selection; CD14+ cells were harvested at 3, 4, 5, 6, 7 and 14 dpi and CD34+ HPCs were harvested at 4 dpi.

Lytic infection was carried out on primary fibroblasts and monocyte-derived macrophages obtained by growing CD14+ monocytes in 50ng/ml PMA containing media for 2 days. For reactivation assays, infected monocytes were differentiated into dendritic cells (DCs) at 3 dpi by incubation with granulocyte-macrophage CSF and interleukin-4 (Peprotech) at 1,000 U/ml for 5 days, followed by stimulation with 500 ng/ml of LPS (Sigma) for 48 hours (as previously described in (64)). Release of infectious virions was assayed by co-culturing of 100,000 differentiated and non-differentiated infected monocytes at the end of the differentiation procedure with HFF cells for 10 days and quantification of GFP positive plaques. Cell number and viability were measured by Trypan blue staining prior to plating.

For UV inactivation, the virus was irradiated in a Stratalinker 1800 (Stratagene) with 200 mjoules.

### Immunofluorescence

Cells were fixed in 4% paraformaldehyde for 10 minutes, permeabilized with 0.1% Triton X-100 in PBS for 10 minutes and blocked in 10% normal goat serum in PBS. Detection of IE-1 was performed by immunostaining with anti-IE1 antibodies (1:100, Abcam ab53495), followed by goat anti-mouse antibody (1:200, AlexaFluor647, Invitrogen A21235) and Hoechst nuclear stain. Cells were visualized in a Zeiss Axioobserver fluorescent microscope.

### qRT-PCR

Total RNA was extracted using Tri-Reagent (Sigma) according to manufacturer’s protocol. cDNA was prepared using qScript cDNA Synthesis Kit (Quanta Biosciences) according to manufacturer’s protocol. Real time PCR was performed using the SYBR Green PCR master-mix (ABI) on a real-time PCR system QuantStudio 12K Flex (ABI) with the following primers (forward, reverse): IE1 (GGTGCTGTGCTGCTATGTCTC, CATGCAGATCTCCTCAATGC) UL138 (GTGTCTTCCCAGTGCAGCTA, GCACGCTGTTTCTCTGGTTA) RNA 2.7 (TCCTACCTACCACGAATCGC, GTTGGGAATCGTCGACTTTG) RNA 4.9 (GTAAGACGGGCAAATACGGT, AGAGAACGATGGAGGACGAC) Anxa 5 (AGTCTGGTCCTGCTTCACCT, CAAGCCTTTCATAGCCTTCC) *Single cell sorting and MARS-seq RNA library construction* Single cell sorting and library preparation were conducted according to the massively parallel single-cell RNA-seq (MARS-seq) protocol, as previously described (45). In brief, cells from latently infected populations of CD14+ monocytes and CD34+ HPCs were FACS sorted into wells of 384 well capture plates containing 2 µl of lysis buffer and reverse transcription (RT) indexed poly(T) primers, thus generating libraries representing the 3’ of mRNA transcripts. Four empty wells were kept in each 384-well plate as a no-cell control during data analysis. Immediately after sorting, each plate was spun down to ensure cell immersion into the lysis solution, snap frozen on dry ice and stored at −80 °C until processed. Barcoded single-cell capture plates were prepared with a Bravo automated liquid handling platform (Agilent). For generation of RNA library, mRNA from cells sorted into capture plates was converted into cDNA and pooled using an automated pipeline. The pooled sample was then linearly amplified by T7 in vitro transcription, and the resulting RNA was fragmented and converted into a sequencing-ready library by tagging the samples with pool barcodes and Illumina sequences during ligation, RT, and PCR. Each pool of cells was tested for library quality and concentration was assessed as described earlier (45).

### RNA sequencing of lytic cells

For generation of a reference lytic RNA library used in the single cell experiments, monocyte-derived macrophages or primary fibroblasts were infected with TB40E-GFP virus at MOI of 5 and used for library preparation at 4 dpi. The libraries were generated from a samples of ~10,000 cells according to the MARS-seq protocol (45).

The lytic fibroblasts derived RNA-seq libraries used as reference in analysis of the natural samples were previously described (14).

### Single cell library construction using 10x platform

Cell suspensions at a density of 700 cells/μl in PBS + 0.04% BSA were prepared for single cell sequencing using the Chromium Single Cell 3’ Reagent Version 2 Kit and Chromium Controller (10x Genomics, CA, USA) as previously described (65). Briefly, 9,000 cells per reaction were loaded for Gel Bead-in-Emulsion (GEM) generation and barcoding. GEM-RT, post GEM-RT cleanup and cDNA amplification were performed to isolate and amplify cDNA for library construction. Libraries were constructed using the Chromium Single Cell 3’ Reagent Kit (10x Genomics, CA, USA) according to manufacturer’s protocol. Library quality and concentration were assessed according to manufacturer’s instructions.

### Sequencing

RNA-Seq libraries (pooled at equimolar concentration) were sequenced using NextSeq 500 (Illumina), at median sequencing depth of ~45,000 reads per cell for MARS-Seq and ~32,000 reads per cell for 10x. Read parameters were: Read1: 72 cycles and Read2: 15 cycles, for MARS-seq and Read1: 26 cycles, Index1: 8 cycles, and Read2: 58 cycles for 10x.

### MARS-seq CD14+ analysis

The analysis of the MARS-seq data was done with the tools described in (45) and (46). The reference was created from the hg19 and TB40E (NCBI EF999921.1) strain of HCMV. The transcription units of the virus were based on NCBI annotations, with some changes based on the alignment results. This includes merging together several transcripts (taking into account that the library maps only the 3’ of transcripts), and adding some antisense transcripts. Reads assignment to wells was based on the batch barcode (4bps) and the well barcode (7bp), and removing reads with low quality of the barcodes. The read itself (37bp) was aligned to the reference using bowtie2 (Langmead and Salzberg, 2012), and the counting of the reads per gene is done based on unique molecular identifiers (UMIs) (8bp). For each batch the leakage noise level was estimated by comparing the number of UMIs in the 2 empty wells, to the total number of UMIs in the batch. Batches with high noise level (>8%) were discarded. Wells with number of reads < 1000 were discarded. The number of wells that were used for further analysis is 3655. Genes with low total number of reads (< 10), with low variability (variance / mean < 1.1), and also ribosomal protein and histones were excluded. By using a multiplicative probabilistic model, and expectation-maximization like optimization procedure, the 3,655 cells were clustered to 6 clusters. The model includes a regularization parameter (=0.5) simulating additional uniform reads to all genes. The clusters are ordered according to the viral content from high to low.

When analyzing correlation in gene expression the error bars represent 95% confidence interval that were calculated by 10,000 bootstrap iteration of the cells in each one of the clusters. The t-SNE plot of the MARS-seq CD14+ cells was calculated with the R package (67), after down sampling each cell to 1000 UMIs.

To exclude background noise, in each one of the batches, all cells with number of viral reads below 3 times the estimated noise at this batch, were excluded.

To estimate the p-value of getting number of reads n, in cluster B, under the null hypothesis of same expression program as in cluster A, a semi parametric bootstrap method was used. First the probability of sampling UMIs for each viral gene was calculated according to the gene expression in cluster A. Then each bootstrap simulation consists of a parametric step and a-parametric step. The parametric step is, for each cell in cluster B, to sample number of UMIs according to the actual number of read in this cell, with distribution over the genes according to the probabilities calculated from cluster A. Then the a-parametric step is a usual bootstrap sampling of the cells in cluster B, and calculate the total number of reads in this cluster B. After doing this simulation 1,000 times, for each viral gene, the mean and the standard deviation of the number of reads in cluster B, under the null hypothesis was calculated. Based on this value, the Z-score of the actual value n was calculated, and a p-value was calculated assuming normal distribution of the number of reads under the null hypothesis. Lastly, these p-values were adjusted for multiple testing, and just the genes with false discovery rate (FDR) of < 0.01 are reported in tables S4 and S5.

### GTEx and GEO analysis

All RNA-Seq, paired end GTEx samples available on July 2016 were used for the analysis. The reference genome that was used was based on hg19 and Merlin strain of HCMV (NC_006273.2). Bowtie2 (Langmead and Salzberg, 2012) was used for alignment with the default parameters, besides the additional flag --local. Pairs with mapping quality less than 30 were excluded. Pairs with only one read aligned to the Merlin sequence were excluded. For each sample possible PCR duplications were removed. The counting of the alignments to the genes was done with HTSeq-count (68). Annotation of gff files is based on NCBI data, with some adjustment taking into account correcting for the non-stranded library. The clustering and Fig. 5B were generated with GENE-E (69). The analysis of the CD34+ GEO samples was carried out in the same way. The list of datasets that were used is presented in Table S2.

### 10X CD34+ Data analysis

We used CellRanger v2.0.0 (70) software with the default settings to process the FASTQ files. The reference was created with the mkref CellRanger command, based on the CellRanger human hg19 reference, and TB40E (NCBI EF999921.1) as was used in the analysis of the MARS-seq data. The de-multiplexing of the Illumina files, and the analysis done with the CellRanger commands mkfastq and count respectively, The raw reads data was extracted with the CellRanger R Kit (70). The t-SNE plot is based on the coordinates calculated by the count command.

### Ethical statement

All fresh peripheral blood samples were obtained after approval of protocols by the Weizmann Institutional Review Board (IRB application 92-1) and umbilical cord blood of anonymous healthy donors was obtained in accordance with local Helsinki committee approval (#RMB-0452-15). Informed written consent was obtained from all volunteers and all experiments were carried out in accordance with the approved guidelines.

### Data availability

All next generation sequencing data files were deposited on Gene Expression Omnibus and will be publically available upon manuscript publication.

---

**Figure S1:**
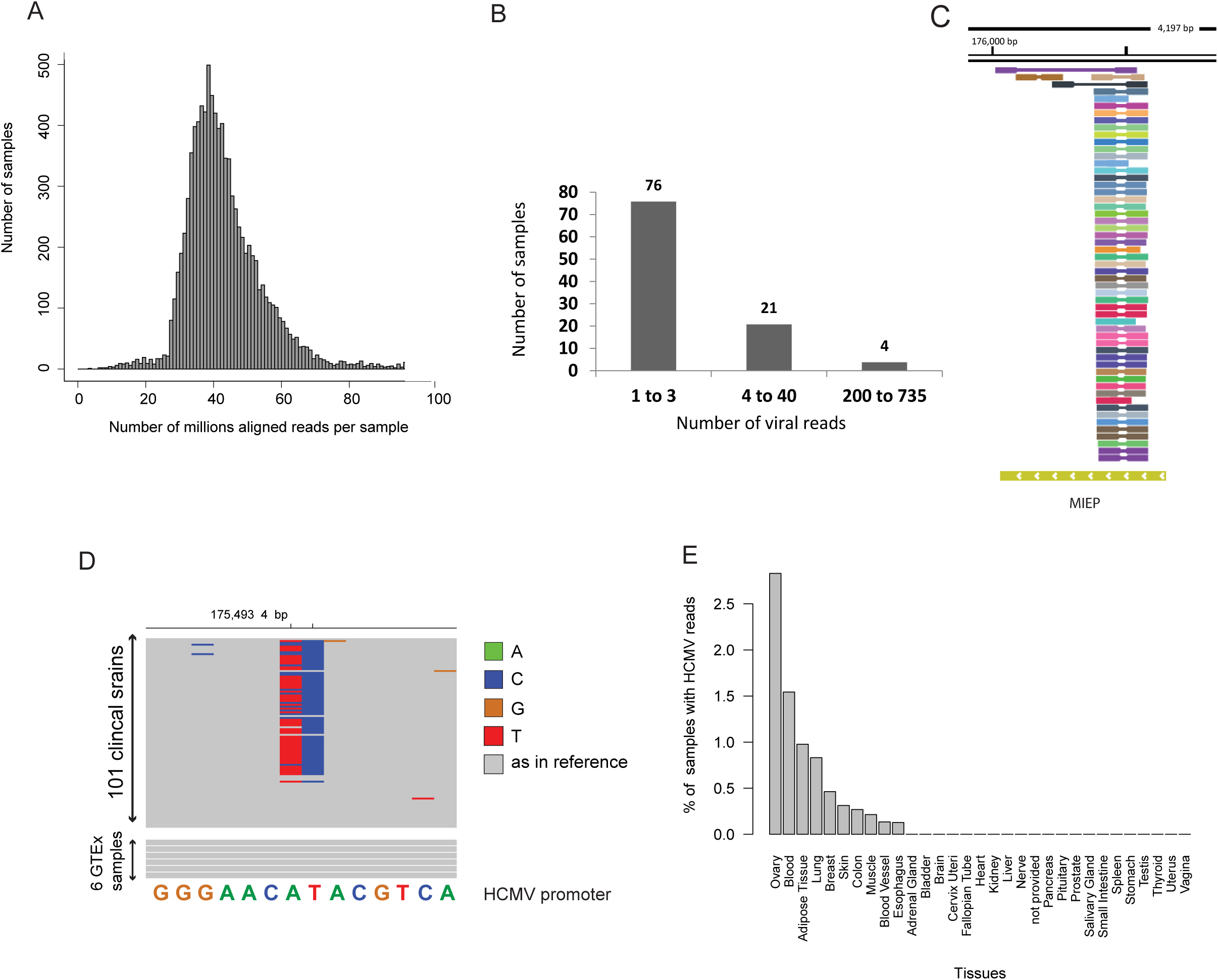
Detection of HCMV reads in natural samples. (A) Distribution of the number of total aligned reads per sample in samples from the GTEx dataset. (B) Distribution of the number of HCMV aligned reads per sample in positive samples from the GTEx dataset. (C) RNA-seq reads from GTEx samples aligned to the MIEP region of HCMV genome colored by sample. (D) Alignment of RNA-seq reads from GTEx samples and sequences of 101 clinical isolates to the MIEP region (positions 175,493 and 175,494 in the viral genome). Base variation from the reference (Merlin strain, which is identical in these sites to the CMV promoter that is used in plasmids) is indicated by a color corresponding to the substituting base. Color legend is on the right. (E) Percentage of samples containing HCMV reads in different tissues. Viral reads from samples containing less than 2 viral reads were filtered out.

**Figure S2:**
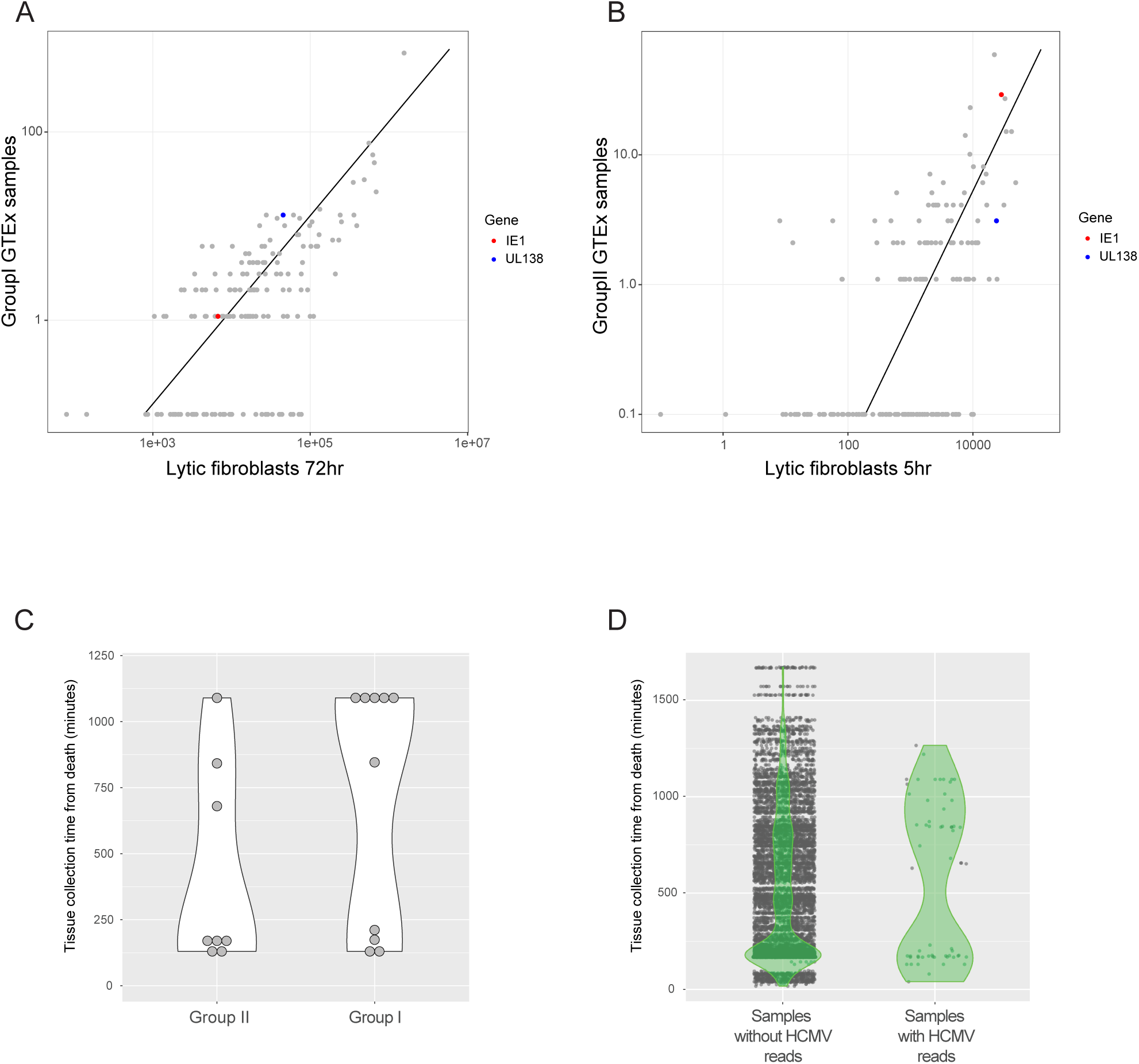
Clustering according to HCMV reads in natural samples. (A) Scatterplot showing read number of viral genes in group I samples from the GTEx database versus lytic fibroblasts 72 hours post infection. (B) Scatterplot showing read number of viral genes in group II samples from the GTEx database versus lytic fibroblasts 5 hours post infection. (C and D) Violin plots showing the time of sample harvesting (measured in minutes after death) versus (C) sample assignment to gene expression group (I or II) and (D) presence or absence of HCMV specific reads in the sample.

**Figure S3:**
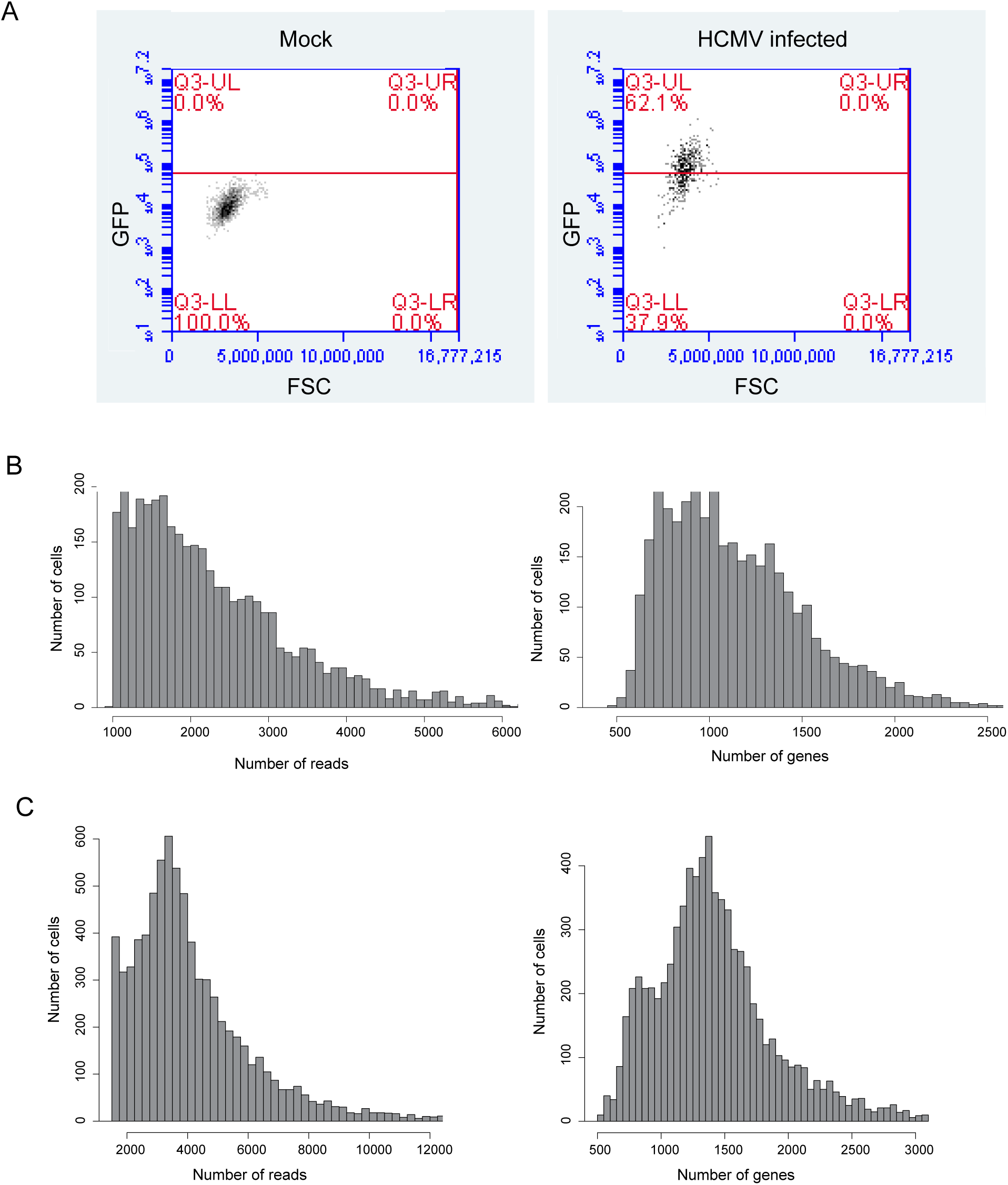
Validation of infection and scRNA library composition. (A) Flow cytometry analysis showing GFP expression level in population of CD14+ monocytes infected with TB40-GFP at 2dpi. (B and C) Bar plots showing distribution of number of reads per cell (left) and number of genes per cell (right) in scRNA-seq data of (B) infected CD14+ monocytes and (C) CD34+ HPCs.

**Figure S4:**
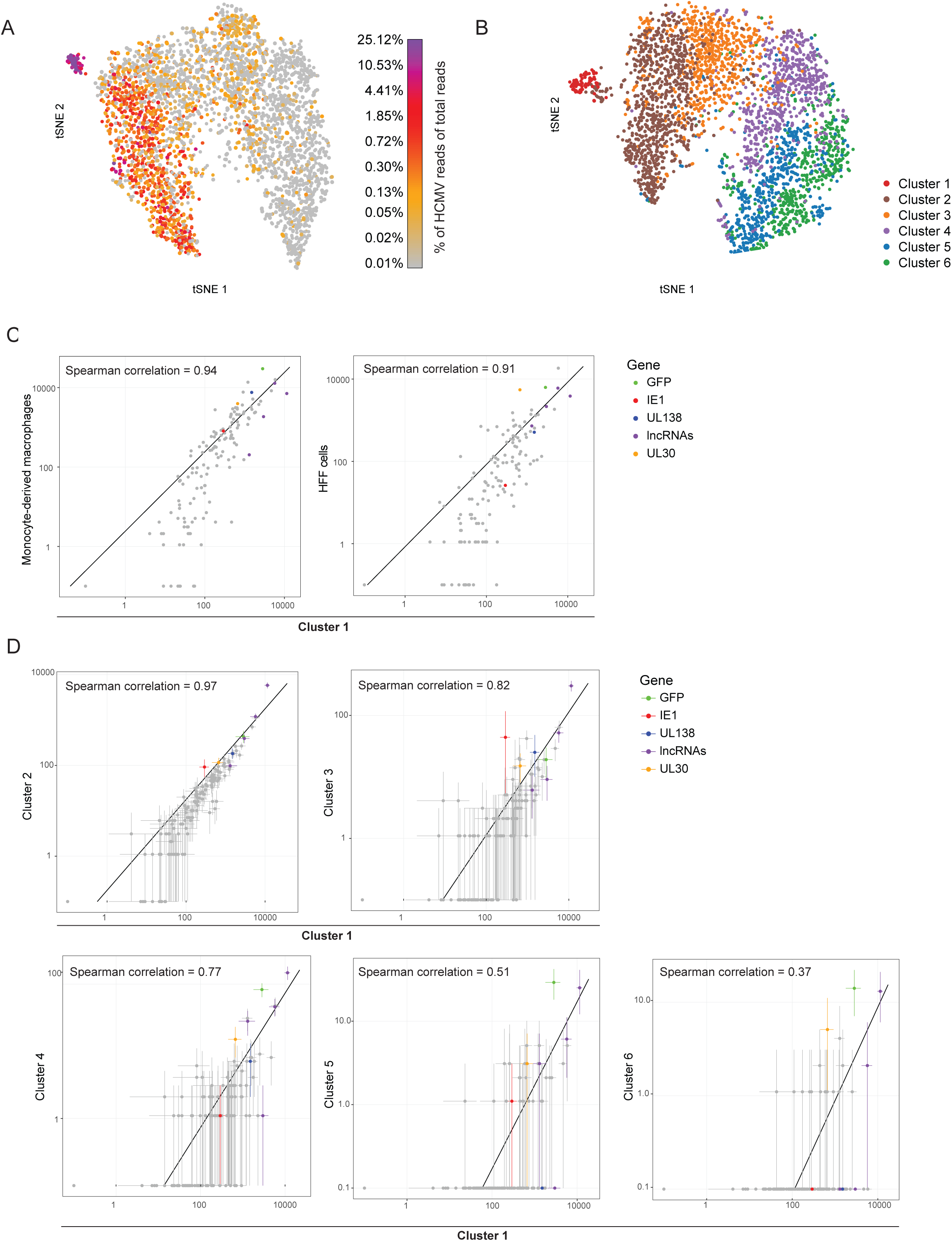
scRNA-seq analysis of latently infected CD14+ monocytes. (A) t-SNE plot of all 3,655 single cells based on host gene expression. Color bar shows the percentage of viral reads from total reads per cell. (B) t-SNE plot of 3,655 single latently infected monocytes based on host and viral gene expression (as shown in Fig. 3A) depicting the separation to 6 clusters as shown in Fig 3B. (C) Scatterplot showing read number of all viral genes in cells from cluster 1 versus lytically infected monocyte derived macrophages at 4 dpi (left panel) or fibroblasts at 3 dpi (right panel). (D) Scatterplot showing read number of all viral genes in cells from clusters 2-6 (labeled on y-axis) versus cells from cluster 1. Horizontal and vertical error bars indicate 95% nonparametric bootstrap confidence interval across cells.

**Figure S5:**
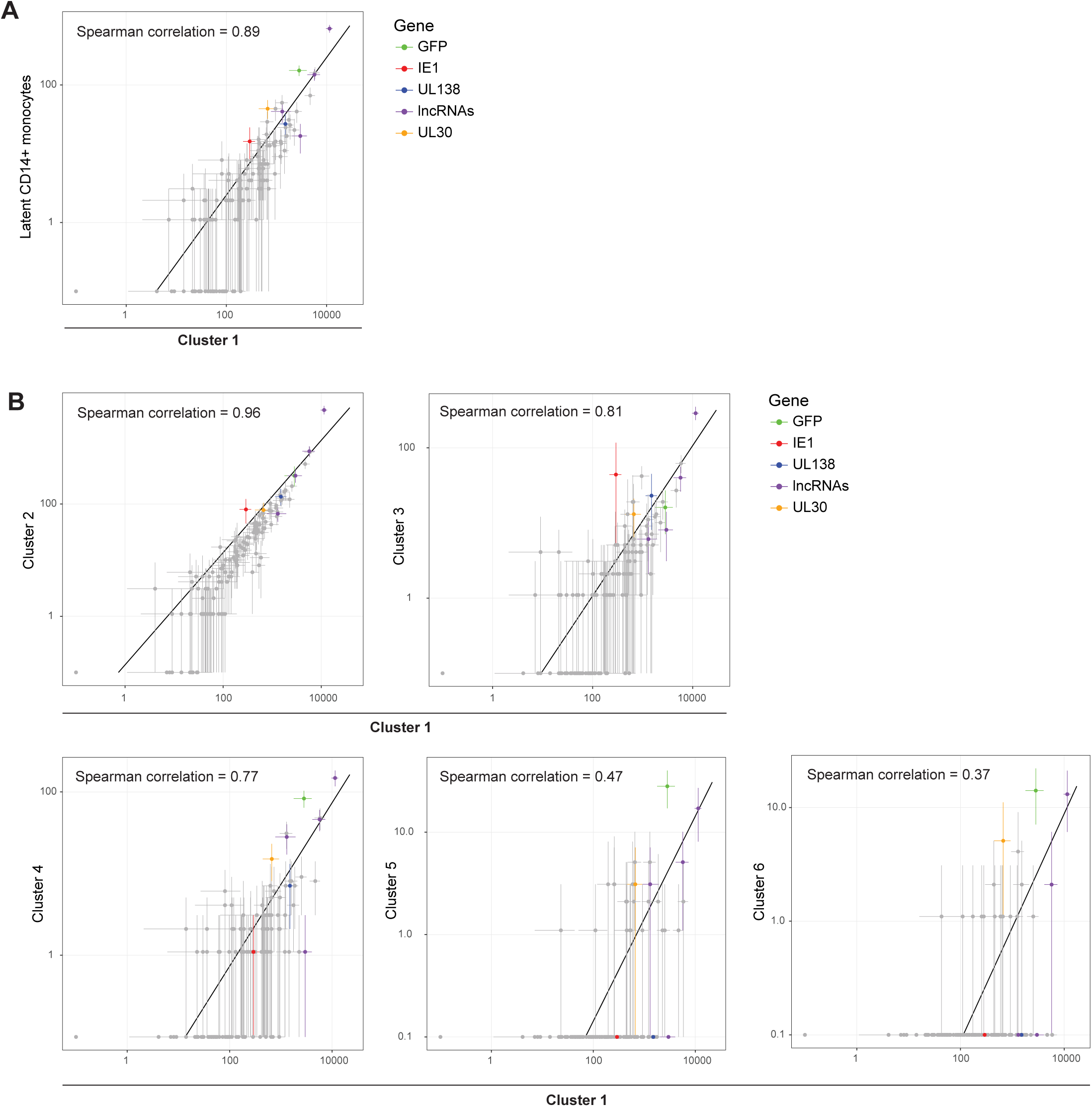
Assessment of lytic noise effect on gene expression in CD14+ scRNA-seq libraries. Scatter plot showing read number of all viral genes in (A) latent cells (defined as cells in which the proportion of viral reads was below 0.5% of total reads) and in (B) cells from clusters 2-6 (labeled on y-axis) versus cells from cluster 1. Analysis was done after exclusion of cells in which viral read counts were lower than the noise cut-off level (See materials and methods). Horizontal and vertical error bars indicate 95% non-parametric bootstrap confidence interval across cells.

